# Stiffening of prostate cancer cells driven by actin filaments – microtubules crosstalk confers resistance to microtubule-targeting drugs

**DOI:** 10.1101/2020.06.14.146696

**Authors:** Andrzej Kubiak, Matteo Chighizola, Carsten Schulte, Natalia Bryniarska, Julita Wesołowska, Maciej Pudełek, Damian Ryszawy, Agnieszka Basta-Kaim, Piotr Laidler, Alessandro Podesta, Malgorzata Lekka

## Abstract

The crucial role of microtubules in the mitotic-related segregation of chromosomes makes them an excellent target for anticancer microtubule targeting drugs (MTDs) such as vinflunine, colchicine, and docetaxel. MTDs affect mitosis by directly perturbing the structural organization of microtubules. By a direct assessment of the biomechanical properties of prostate cancer cells exposed to different MTDs using atomic force microscopy, we show that cell stiffening is a candidate mechanism through which cancer cells preserve the original phenotype in response to the application of MTDs. While changes in cellular rigidity are typically mainly attributed to remodeling of the actin filaments in the cytoskeleton, here we provide evidence that cell stiffening can be driven by a crosstalk between actin filaments and microtubules in drug-treated cells. Our findings improve the interpretation of biomechanical data obtained for living cells in studies of various physiological and pathological processes.

## Introduction

Microtubules (MTs) are filamentous structures constituting one of the major components of the cell cytoskeleton. They are cylinders composed most often of 13 longitudinally arranged protofilaments. Individual protofilaments are composed of tubulin heterodimers built up from α- and β-tubulin linked together by non-covalent bonds forming α/β-tubulin heterodimer^1^. Inside the cell, microtubules start from a microtubule-organizing center (MTOC) and span the entire cytoplasm, ending at a cell membrane. MTs provide functional pathways for the transport of cellular cargo and they are involved in intracellular signaling^1^. The most important role of microtubules is the participation in mitosis. During this process, they form a mitotic spindle used by cells to separate replicated chromosomes into two newly created nuclei^2,3^. Thus, any agents affecting microtubule dynamics will introduce severe consequences for cell division by disabling the completion of mitosis, and thereby, by an inhibition of cell proliferation. Such effect has been in the center of attention of clinical oncologists for many years, leading to the use of taxanes and vinca alkaloids in anticancer therapy of various cancers including lung, breast, prostate, gastric, esophageal, and bladder, as well as squamous cell carcinoma of the head and neck^4–6^.

Since the invention of atomic force microscopy (AFM^7^) and its first use for characterizations of mechanical properties of cells^8–10^ the spectrum of possible AFM applications has widened enormously. A variety of AFM-based mechanical measurements provide evidence that changes in cell rigidity are related to the cell cytoskeleton, mainly to actin filaments^11,12^. Gathered evidence shows that AFM-based mechanical measurements contribute to a better understanding of physical mechanisms involved in the cellular response to various conditions, including chemical treatments.

Antimitotic agents belonging to microtubule-targeting drugs (MTDs) such as docetaxel improve prostate cancer patient’s survival^13^ but, during treatment, cancer cells often develop drug resistance. Despite the known antitumor effect, still it is not known why some cells are resistant to the MTDs. Multidrug resistance linked with overexpression of ABC transporters is one of the possible explanations. Another possible explanation is changes in the expression of tubulin isotypes, which prevent interaction between the drug and its target^14^. Mechanisms of MTDs binding to α/β tubulin heterodimers affect their dynamics by stabilizing or destabilizing microtubular network^15^. Therefore, strong prerequisites indicate that biomechanical cues underlie the cellular response of prostate cancer cells to drugs against microtubules. Molecular mechanisms involved in the binding of MTDs are, theoretically, independent of the cell types indicating that all cancers can be treated with such drugs. However, there exists a group of cancers that do not respond positively to such a treatment, showing cells resistant to the drug treatment, as in the case of prostate cancers. Here, we combine AFM and cell biological methods to study changes in the mechanical properties of prostate DU145 cancer cells treated with three MTDs drugs i.e., vinflunine (VFL), docetaxel (DTX), and colchicine (COL) at concentrations below *IC*_*50*_. We identified the cell stiffening as a process through which cancer cells preserve the original phenotype in response to applied drugs.

## Results

### Viability study shows that attached DU145 cells are more resistant to low MTDs doses

We started our analysis with the evaluation of MTDs’ effect on the viability and proliferation of DU145 cells (**Fig. 1a)**.

**Figure 1.**
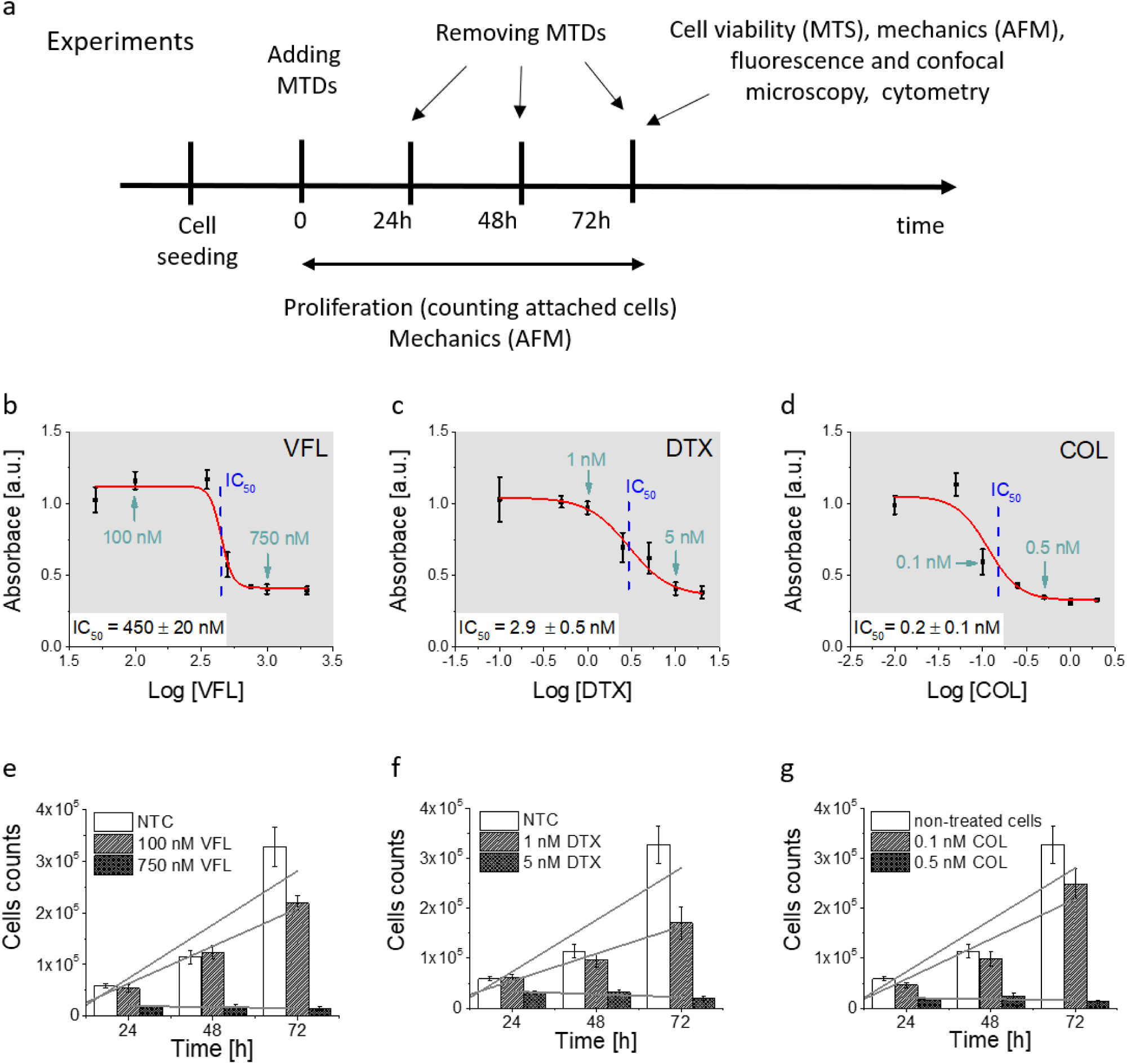
Prostate DU145 cells respond differently to distinct MTDs drug concentrations at different time-points. **a**, Illustration of experimental steps and methods applied. **b–c**, Dose-response curves for cells exposed to VFL (**b**), DTX (**c**), and COL (**d**). Calculated from the fit, IC_50_ were 450 ± 20 nM (VFL), 2.9 ± 0.5 nM (DTX) and 0.2 ± 0.1 (COL). **e-g**, Proliferation level of prostate DU145 cells determined for low and high doses for VFL (**e**), DTX (**f**), and COL (**g**). A slope of linear fits determines a rate of proliferation. NTC cells denote non-treated DU145 cancer cells. Data are shown as mean ± standard deviation (s.d.) from at least n = 3 measurements; IC_50_ is expressed as a fitted value ± standard error, (s.e.). Arrows indicate the position of low and high concentrations of drugs chosen for further experiments.

To verify how low and high MTDs concentrations deviate from the *IC*_*50*_, the MTS assay was applied to cells exposed to different drug concentrations, i.e. 50 nM – 2000 nM for VFL, 0.1 nM to 20 nM for DTX, and 0.01 nN to 2 nM for COL. MTS assay quantifies the fraction of metabolically active cells (through dehydrogenase activity). Recorded absorbance was normalized to a viability level of NTC cells and plotted against MTDs drug concentration in the logarithmic scale (**Fig.1 b-d**). *IC*_*50*_, obtained from the fit of a dose-response function, was 450 ± 20 nM (VFL, **Fig. 1b**), 2.9 ± 0.5 nM (DTX, **Fig. 1c**), and 0.2 ± 0.1 nM (COL, **Fig. 1d**), respectively.

The proliferation rate was assessed by counting cells attached to the glass surface after their incubation with MTDs for 24h, 48h, and 72h. Results show a strong decrease in the cell number for high drug concentrations i.e., for 750 nM for VFL, 5 nM DTX and 0.5 nM COL (**Fig. 1 e-g**), regardless of the culture time. The number of cells drops to the level of 10% – 20% with respect to the number of alive, non-treated cancer (NTC) cells. For low MTDs concentrations, i.e. 100 nM for VFL, 1 nM for DTX, and 0.1 nM COL, the number of cells after 24 h and 48 h of culture remained at the same level as for NTC cells regardless of the drug type. The decrease in the number of cells was observed only after 72 hours. A drop by 33%, 48%, and 23% for VFL, DTX, and COL, respectively, was accompanied by the smallest proliferation rate **(Suppl. Note 1)**.

In our study, low MTDs concentrations are placed below *IC*_*50*_ while high concentrations are above *IC*_*50*_. 72h exposure of DU145 cells to low concentrations of MTDs reveals a cell population that contains a strongly dominating fraction of proliferating cells, while at high concentrations many of the cells died through both apoptosis and necrosis. All cells heavily affected by MTDs are detached from the surface. This is the main reason why the *IC*_*50*_ (MTS, bulk measurements) does not fully correlate with the decrease in the number of cells attached to the surface (proliferation). Therefore, our results show that DU145 prostate cells seem to be more resistant to COL, than to VFL and DTX.

### NTC and VFL-treated cells show a similar level of calcein efflux

To elaborate on how the viability of cells is linked with the capability of DU145 cells to release MTDs, in our next step, we decided to trace acetoxymethyl (AM) ester derivative of calcein using flow cytometry in NTC and MTDs treated cells **(Fig. 2)**. Calcein-AM is a non-fluorescent, uncharged molecule that can freely permeate the plasma membrane of viable cells. Inside the cell, it is converted to a fluorescent form, calcein, upon cleavage of the lipophilic blocking groups by nonspecific esterase in the cytoplasm^16,17^. Cytometric study shows that within the analyzed population of DU145 cells (NTC, 5.6%, **Fig. 2a**), there are minoritarian groups of cells actively pumping calcein out, i.e. side population. The side population denotes the shift of fluorescence proportional to the loss of calcein pumped out or accumulated inside the cells^17^. The size of this group decreases for cells treated with MTDs. The largest drop, about two-fold as compared to non-treated DU145 cells, was observed for COL (2.7%, **Fig. 2d**) and DTX (3.6%, **Fig. 2c**) treated cells. In the case of VFL treatment, the population of these cells was only slightly smaller (4.9%, **Fig. 2b**).

**Figure 2.**
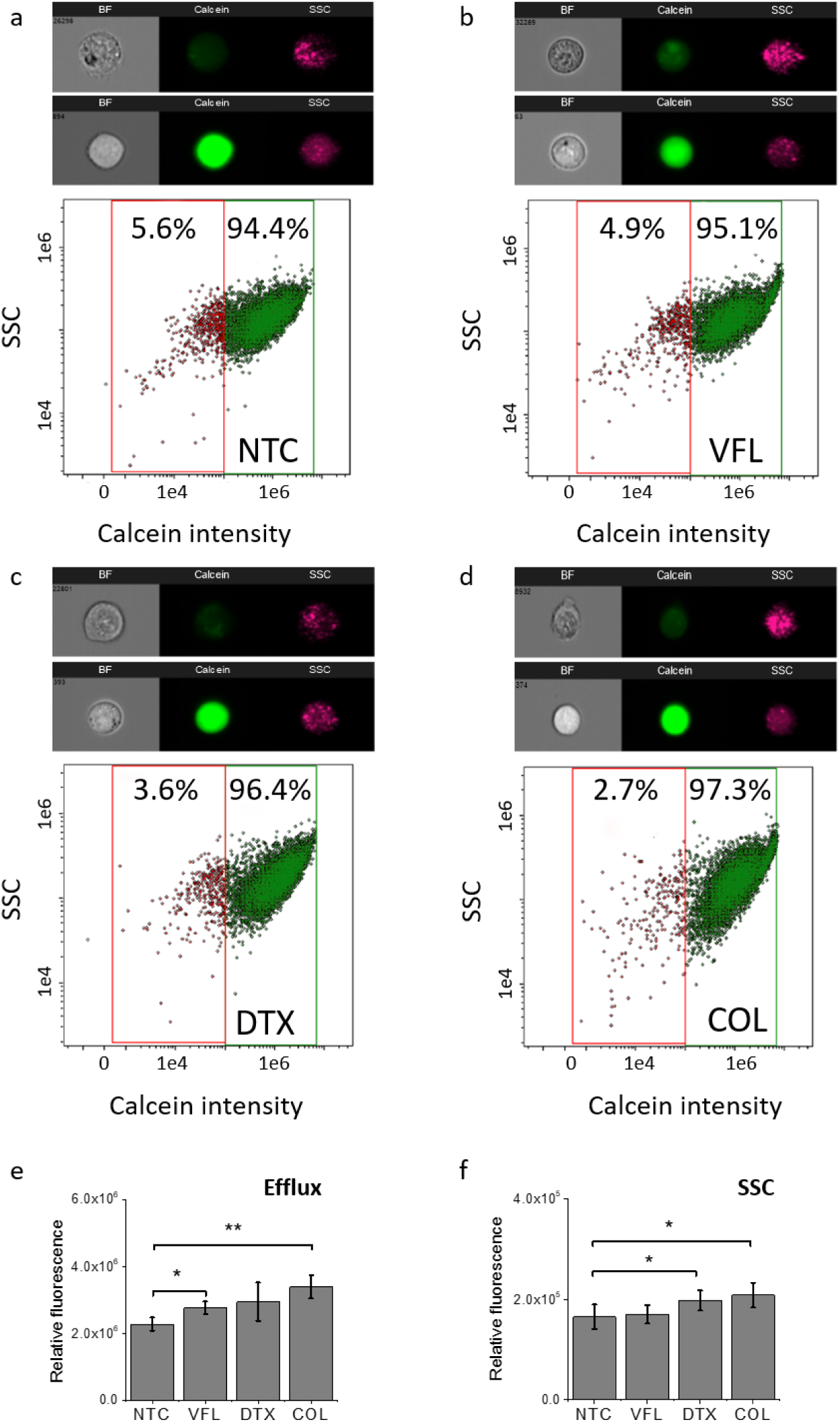
Calcein fluorescence intensity changes with the MTDs type. **(a - d)** Correlative dot plots of side scatter light (SSC) and calcein intensity within the cell population 72h after MTDs treatment. The high SSC/high calcein intensity subpopulation was observed after cell treatments with VFL and COL. ImageStream obtained cell images are representative of the cells gated in NTC (green) and MTDs-treated (magenta) cell populations. (**e-f**) Average fluorescent intensity of calcein calculated as a mean ± s.d. (n = 3 independent measurements). Statistical significance was estimated by unpaired t-Student test at the level of 0.05.

Together with the decreasing number of cells and the pumping of calcein to the extracellular microenvironment, a shift of a cytometric population towards higher calcein intensity appears. The observed MTDs-related retention of calcein in treated DU145 cells could be indirectly attributed to drug accumulation (**Fig. 2e&f)**.

### Microtubule rearrangements depend on MTDs binding site

MTDs such as vinflunine, docetaxel, or colchicine are known to bind at distinct locations in the *α/β* tubulin heterodimer (**Fig. 3a)** and affect the organization of the microtubular network. Thus, we visualized the organization of the microtubules inside the DU45 cells after 72 hours of incubation with a specific drug at low and high concentrations (**Fig. 3b-h)**.

**Figure 3.**
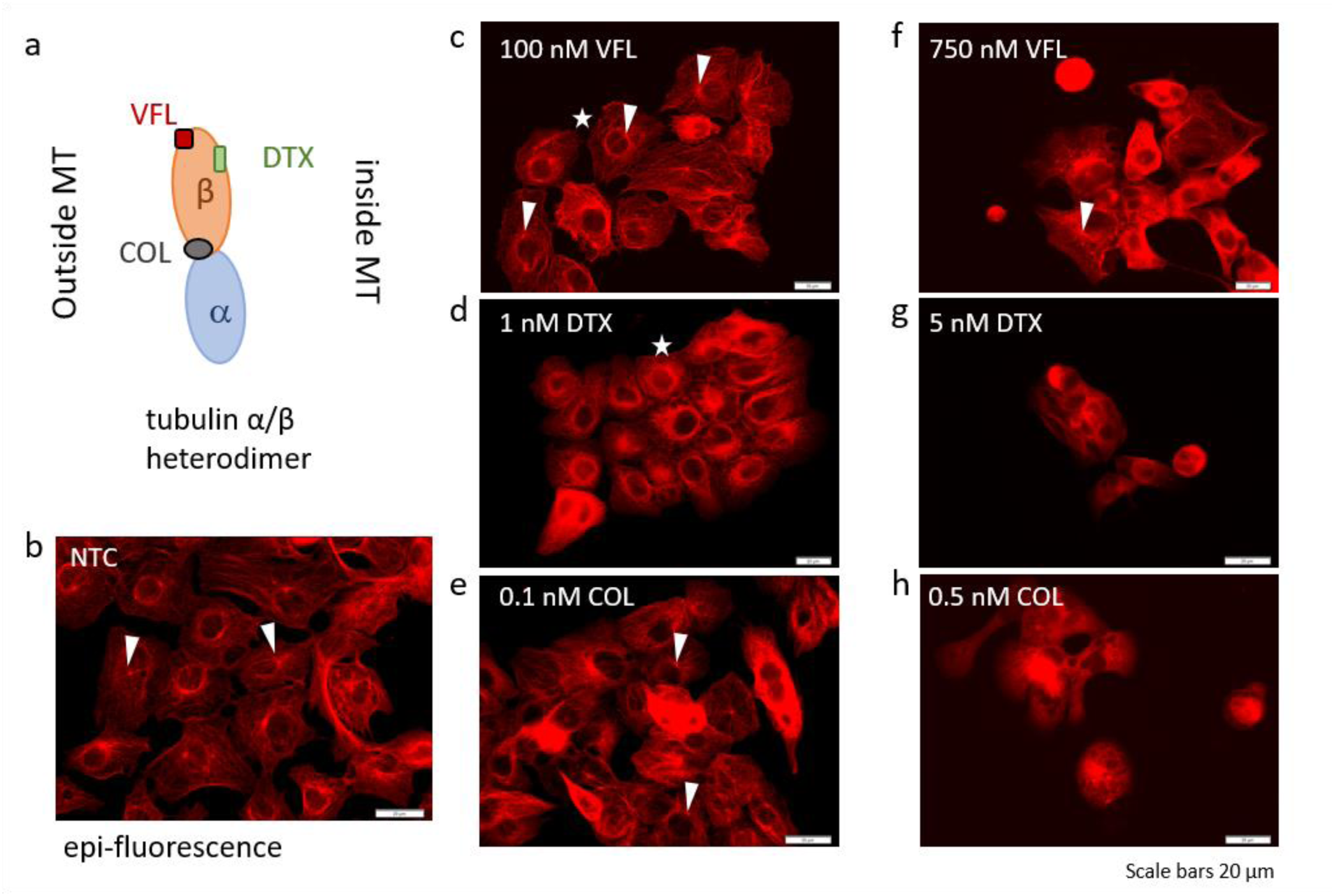
MTDs bind to distinct sites located at the α/β tubulin heterodimer affecting differently the organization of the microtubular network. **a**, A scheme illustrating the binding of VFL, DTX, and COL to α/β tubulin heterodimer. **b-h**, Epi-fluorescent images showing the organization of microtubules in DU145 cells upon treatment for 72 hours with low and high MTDs doses (white arrows show MTOCs while stars indicate MT rings formed around cell nuclei).

In NTC cells, microtubules exhibit multi-branched tubulin networks with clearly visible MTOCs located close to the cell nucleus (indicated by white arrows, **Fig 3b**). Microtubules alone form long fibers that span from MTOC towards the cell membrane. For cells treated with low MTDs concentrations, the degree of microtubule re-organization is a drug-dependent (**Fig. 3c-e**). In cells treated with 100 nM VFL, in which MTOCs are visible, the microtubular network is weakly affected (**Fig. 3c)**. Here, proliferating cells are still present (**Fig. 1, Suppl. Fig. S7**).

Around cell nuclei, MTs form a ring, which is strongly or weakly visible under fluorescence (marked by stars in e.g. **Fig. 3c**). The presence of both MTOCs and rings indicate that, after 72h of culture in the presence of MTDs, most of DU145 cells were not at all or weakly affected by these drugs. The microtubular network was strongly preserved for VFL-treated cells (**Fig. 3c**). In the case of prostate cancer cells exposed to 1 nM DTX for 72h, MTOCs became less visible and the microtubular network seems to shrink (**Fig. 3d**). This was accompanied by the aggregation of microtubules in a form of thicker rings surrounding the cell nucleus. Altered organization of MTs is associated with a lower proliferation rate (**Suppl. Note 1**) and cell viability level (**Fig. 1)**. The effect of 0.1 nM COL on DU145 cells also induced a rearrangement of the microtubule network (**Fig. 3e)**, similar to that obtained for 100 nM VFL. Here, MTOCs were visible only in some cells. These changes in the MT network can be rated as follows (from the strongest to weakest): DTX > COL > VFL. This agrees with the MTS assay, from which one can expect large changes in cells treated with DTX, COL, and VFL (**Fig. 1**).

For high MTDs concentrations, the clustering of cells was recorded (**Fig. 3f-h)**. Although many cells were affected, still single cells were detected that showed organized MTOCs (e.g. **Fig. 3f**). But in the majority of cells, the microtubular network was fully disorganized (disassembled), especially in the case of DTX and COL, for which a destruction of MTs is expected as it could be linked with apoptosis^18^. Changes in the microtubular network were accompanied by morphological alterations observed for cell nuclei **(Suppl. Note 2, Suppl. Fig. S1; Suppl. Fig. S2)**. High MTDs doses cause a fragmentation of nuclei, especially for DTX and COL, while low MTDs doses reveal a strong decrease of nuclei surface area circularity and roundness for DTX treated cells **(Suppl. Fig. S1c-e)**. The results obtained for low doses confirm the postulate that drugs affecting microtubules at small concentrations are not sufficient to destroy them, but they tend to change MTs dynamics^6^.

### DU145 cells stiffen only when treated with VFL

Changes in MT organization roughly correlate with the results of cell viability. At the same time, according to already published results^19–21^, MT organization also is expected to affect the mechanical properties of cells. To investigate the role of biomechanics in drug resistance of cancer cells, we employed AFM to characterize the mechanical properties of treated cells^22,23^. In particular, we used AFM to obtain an overview of the temporal evolution of cellular rigidity upon exposure to different concentrations of MTDs. The relative change *ΔE*_*rel*_ of Young’s modulus *E*, defined as *ΔE*_*rel*_ = *(E*_*drug*_ *– E*_*NTC*_*)/E*_*NTC*_, is quantified using Hertz contact mechanics (**Materials and Methods, Suppl. Table S1, Suppl. Note 3; Supp. Fig. S2**).

We first carried out a characterization of the relative changes in the mechanical properties of cells, measured within cell center i.e. above the cell nucleus at different time-points, i.e, after 24h, 48h, and 72h of culture in the presence of MTDs. The results show that MTDs type, dose, and time alter the biomechanical properties of prostate DU145 cells (**Fig.4a-c, Suppl. Fig. S3, Suppl. Table S1**).

**Figure 4.**
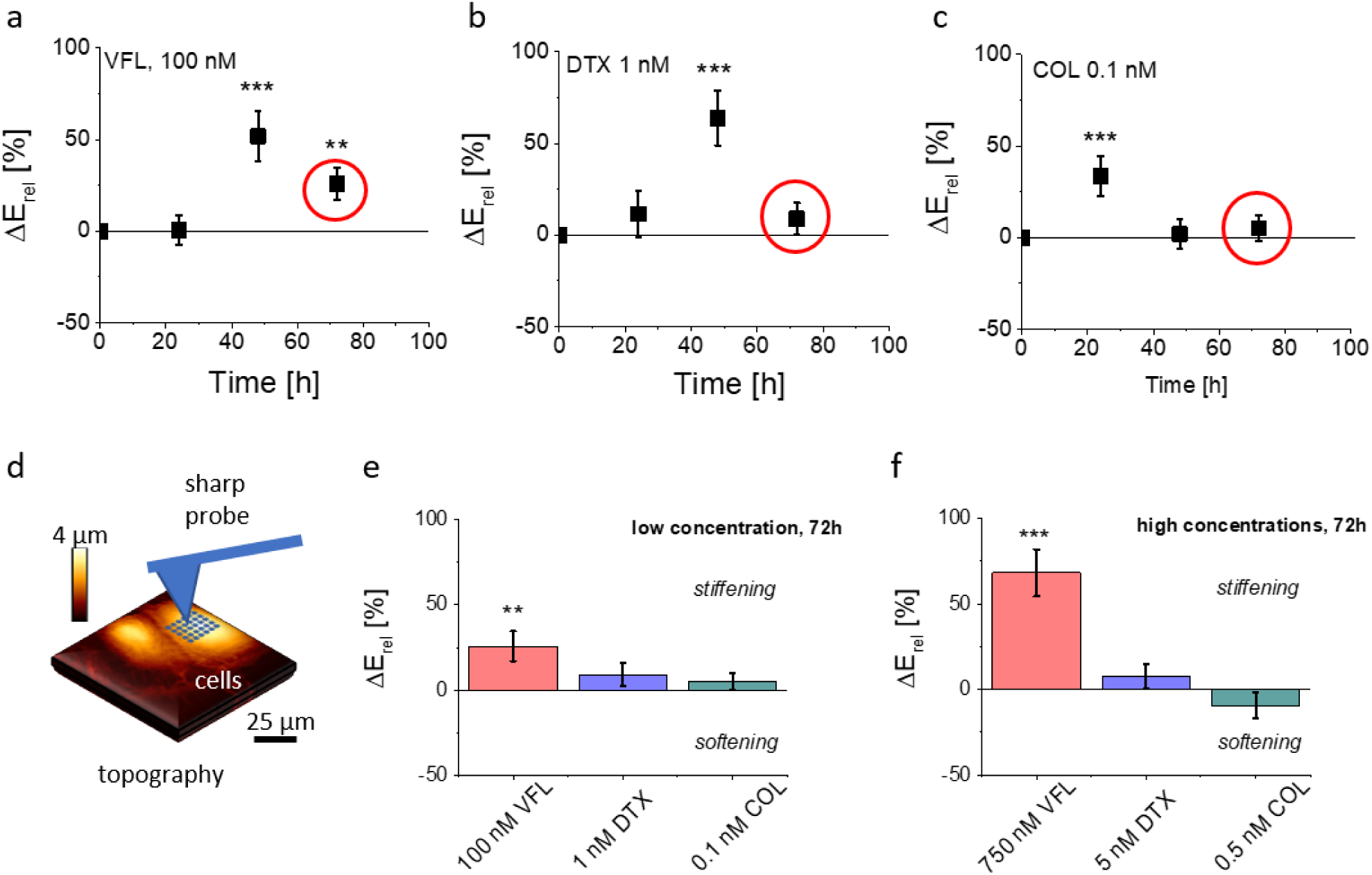
Mechanical properties of DU145 cells depend on the dose and MTDs type. **(a-c)** Relative Young’s modulus change, ΔE, determined for cells treated with low MTDs doses in relation to NTC cells after 24h, 48h, and 72h of the incubation with the corresponding MT drug. Depending on the MTDs, a maximal increase of cell deformability was observed. Circle indicates values, for which a decrease of cell proliferation was observed (**Fig. 1**). (**d**) Measurements were conducted using AFM in indentation mode using pyramidal probes. (**e-f**) The 72h treatment of DU145 cells exposed to MTDs shows that a large change in Young’s modulus for cells treated with VFL, regardless of the applied drug concentration, indicating cell stiffening. DTX action leads to weak stiffening regardless of the applied drug concentration while COL treatment to either stiffening (**e**, high drug doses) or softening (**f**, low drug dose) of DU145 cells. All data were normalized by the Young’s modulus of NTCs measured at the same time-point. Statistical significance was estimated by unpaired t-Student test at the level of 0.05 (** p < 0.01, *** p < 0.001.

For each MTD, a maximum of cell rigidity was observed. For VFL and DTX after 48h of culture with the drug, while for COL the maximum occurred after 24h of culture. Interestingly, the maximum rigidity is not related to the proliferation rate (**Fig. 1)**, showing the decrease in the number of cells after 72h of drug exposure. Also, such an increase was weakly associated with the organization of the microtubular network as MTs re-arranged strongly after 72h of culture with MTDs. Already published data indicate that for high concentrations, microtubule-targeted drugs can be broadly grouped into microtubule-stabilizing agents and destabilizing agents achieved by either promoting microtubule assembly or by triggering their disassembly into tubulin dimers^3^. The consequence is the re-arrangement of microtubules. Here, high MTDs concentrations cause heavy damages to the microtubular network **(Fig. 3f-h)**, for DTX and COL added to cell culture. Observed large effect of MTs damage was not related to relative change in mechanical properties, neither for low nor high MTD concentrations.

Considering the occurrence of maximal cell rigidity, in our study, we focused on cells exposed to low concentration of MTDs after 72h of culture, for which proliferation rate decreased. AFM indentation experiments showed that prostate DU145 cells stiffen (positive *ΔE, p* < *0.01)*) in response to VFL treatment. After the same time, cells treated with DTX and COL revealed changes in mechanical properties that are statistically not significant (*p* > *0.2*). These results are in contradiction with the fluorescence images of microtubular networks, showing that, for proliferating cells, the morphology of 100 nM VFL-treated cells is very similar to that of NTC cells, while the largest morphological changes are observed for 1 nM DTX and 0.1 nM COL– treated cells, for which less obvious changes in cell mechanics are observed.

### Stiffening of VFL-treated cells occurs within the nuclear region

To probe more thoroughly the mechanical changes occurring after 72h treatment with MTDs, we performed combined AFM topographical and mechanical measurements, and analyzed the elastic response of the cells at different indentation depths^23^ (**Fig. 5, Materials and Methods).** Each recorded force curve was linearized, and two linear regions, corresponding to different Hertzian regimes, were typically identified. The first regime (shallow indentation) corresponding to the 5-15% relative indentation range normalized to the local cell height), is expected to represent mainly the mechanical response of the cell membrane coupled with the actin cortical layer; the second regime (deep indentation) corresponding to the 20–40% relative indentation range, is expected to be strongly affected by the response of the nucleus and the surrounding MT network. Given the peculiar organization of the MT network around the nucleus, we defined nuclear masks to focus our attention on the nuclear region of the cells (**Fig. 5b-d**). This methodology enabled us to analyze the deformability of different cellular regions and layers, according to the following classification: whole cell, from cell clusters (as DU145 cells tend to grow as clusters), nuclear region probed at the deep and shallow indentations, and cell periphery probed at shallow indentation (**Fig. 5c-d**). The applied finite thickness correction minimized the effect of the underlying glass substrate and local cell height variations (**Suppl. Notes 4&5; Fig. S4**). Elasticity maps revealed that in the case of shallow indentations, a large increase of Young’s modulus was observed at the cell periphery (**Fig. 5c**), while the nuclear region appeared softer. At deep indentations, the cell periphery appeared to have lower Young’s modulus as compared to the nuclear regions (**Fig. 5d**).

**Figure 5.**
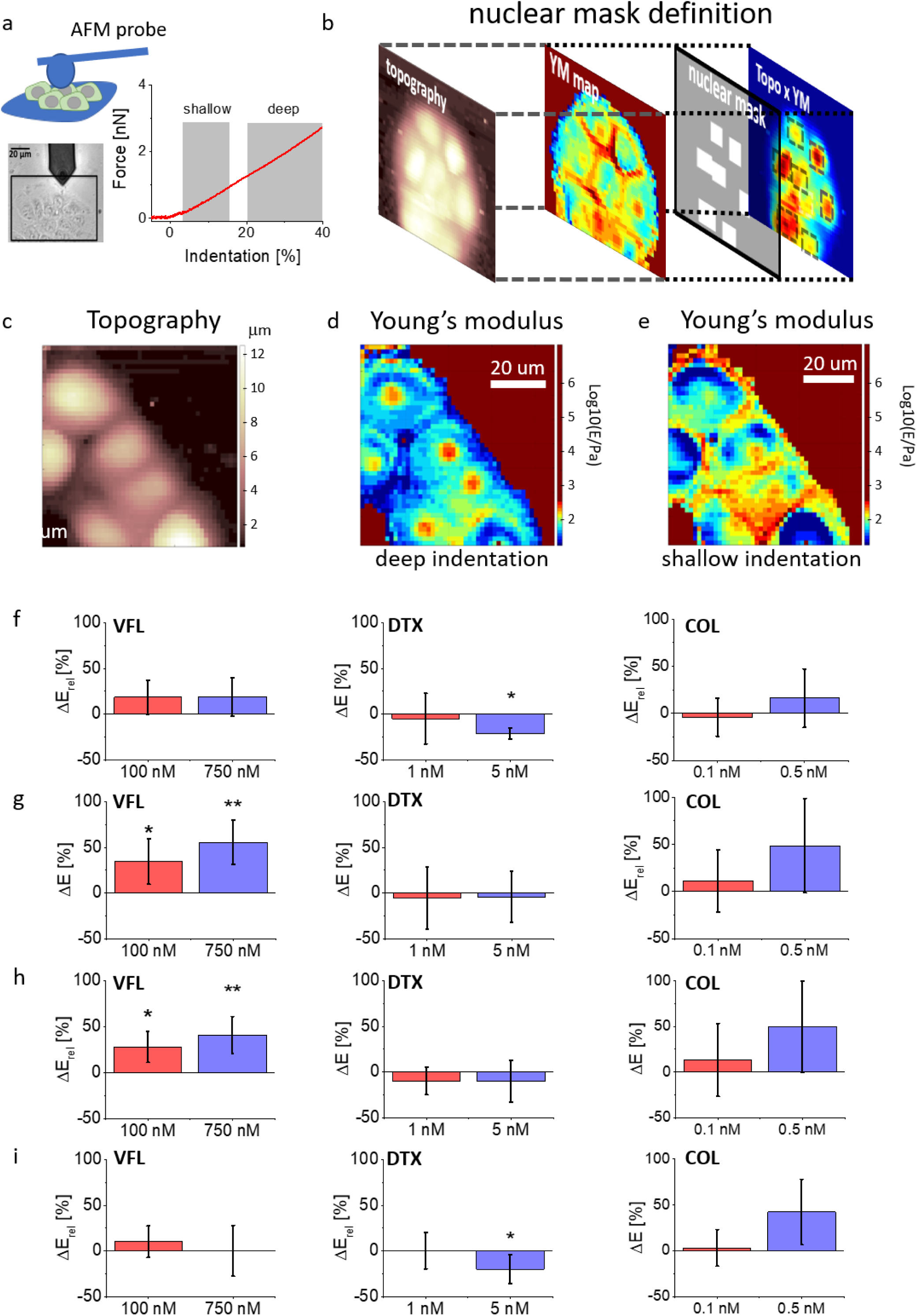
Multicomponent mechanical analysis of cell clusters. **(a)** Exemplary optical image of prostate cancer cell clusters with highlighted regions selected for the indentation measurements (the colloidal probe at the end of the AFM cantilever is visible as a darker circular shadow). Exemplary linearized force curves collected for VFL treated cells with highlighted shallow and deep indentation regions. **(b)** Nuclear mask definition used further to sort the data recorded around cell nuclei. **(c)** Topography of a cell cluster created based on contact point positions. **(d-e)** Young’s modulus maps obtained at **(d)** shallow and **(e)** deep indentations. **(f-i)** Relative Young’s modulus change, ΔE, determined for DU145 cells treated with specific MTDs concentrations for the whole cell (**f**), the nuclear region at shallow **(g)**, and deep **(h)** indentations, and at the cell periphery at shallow indentation (**i)**. Error bars were calculated using the error propagation method. Statistical significance was estimated by unpaired t-Student test at the level of 0.05 (* p < 0.05; ** p < 0.01).

The obtained Young’s moduli for DU145 cells show changes that are strongly dependent on the cellular regions and the used drug (**Fig. 5; Suppl. Table S2**). Mechanical properties of COL-treated cells show a very weak tendency to stiffen, especially for 0.5 nM drug concentration, however, experimental errors are large enough to exclude statistical significance (*p* varies between 0.0584 to 0.8905). This brings the conclusion that colchicine does not affect the mechanical properties of prostate cells. Relative Young’s modulus changes in DTX-treated prostate cancer cells do not reveal any alterations in mechanical properties at low (1 nM) concentration. Changes in cell mechanics were observed for high (5 nM) concentration but instead of stiffening, a softening was noted within a peripheral region. No difference as compared to NTC was observed within a nuclear region at both indentation regimes. DTX-induced softening at the cell periphery (**Fig. 5i**) was dominating in the overall cell softening observed when the whole cells were measured (**Fig. 5f**). In the case of VFL, changes in cell mechanics were more pronounced for nuclear regions at both applied MTD drug concentrations and indentation regions. Lack of *ΔE* variations within the cell periphery and at the level of the whole cell indicates that the mechanical properties measured for individual cells independently of the indentation depth are not affected by changes induced by MTD within nuclear regions.

In summary, DTX and VFL have two different effects on DU145 cells. DTX induces cell softening only at the cell periphery while VFL generates the stiffening of cells within a nuclear region.

### MT network around cell nuclei is not responsible for the stiffening of VFL-treated cells

Changes in mechanical properties of DU145 cells induced by the presence of MTDs are supposed to be associated with the alterations of cytoskeletal networks, especially MTs. Rings observed around cell nuclei in epi-fluorescence images (**Fig. 3c-e)** suggest a higher MT density close to cell nuclei, thus, MT crowding around the nuclei could be responsible for deformability changes induced by MTDs. To test this hypothesis, we employed confocal microscopy to visualize whether the spatial organization of microtubules around the cell nucleus is different in treated and non-treated prostate cancer cells. Typically, the images of the cytoskeleton are recorded close to the surface, to which cells are attached. The AFM measurements characterized also the mechanical properties by applying an external load to the apical region of the cells. Therefore, we also recorded an apical section of the cells. We concentrate on the organization of MTs around cell nuclei in cells treated with low MTDs concentrations. At high MTDs concentration, the organization of the microtubular network is heavily affected and can be seen under epi-fluorescent microscopy (**Fig. 3f-h).** Detailed analysis of the three-dimensional (3D) organization of microtubules in DU145 cells treated with low MTDs doses are included in the **Suppl. Note 6**.

Lack of evident MTs re-organization around the cell nucleus guided us towards considering a crosstalk phenomenon between actin filaments and microtubules^24,25^. Actin filaments play multiple roles regarding the microtubular network by the guidance of filament growth^26^ or crosslinking and anchoring of microtubules within its network^27^. Thus, any disruption of the microtubular network will affect the organization of the actin cytoskeleton and vice versa inducing the chemoresitance of cancer cells^24,28,29^. Thus, we checked whether changes in the mechanical properties of cells might be related to the crosstalk phenomenon by analyzing the organization of actin filaments (**Fig. 6; Suppl. Fig. S5**). After 72 hours of incubation with MTDs, the formation of thick actin bundles was induced in cells. The strongest formation of actin bundles is observed for VFL-treated cells what explains their stiffening.

**Figure 6.**
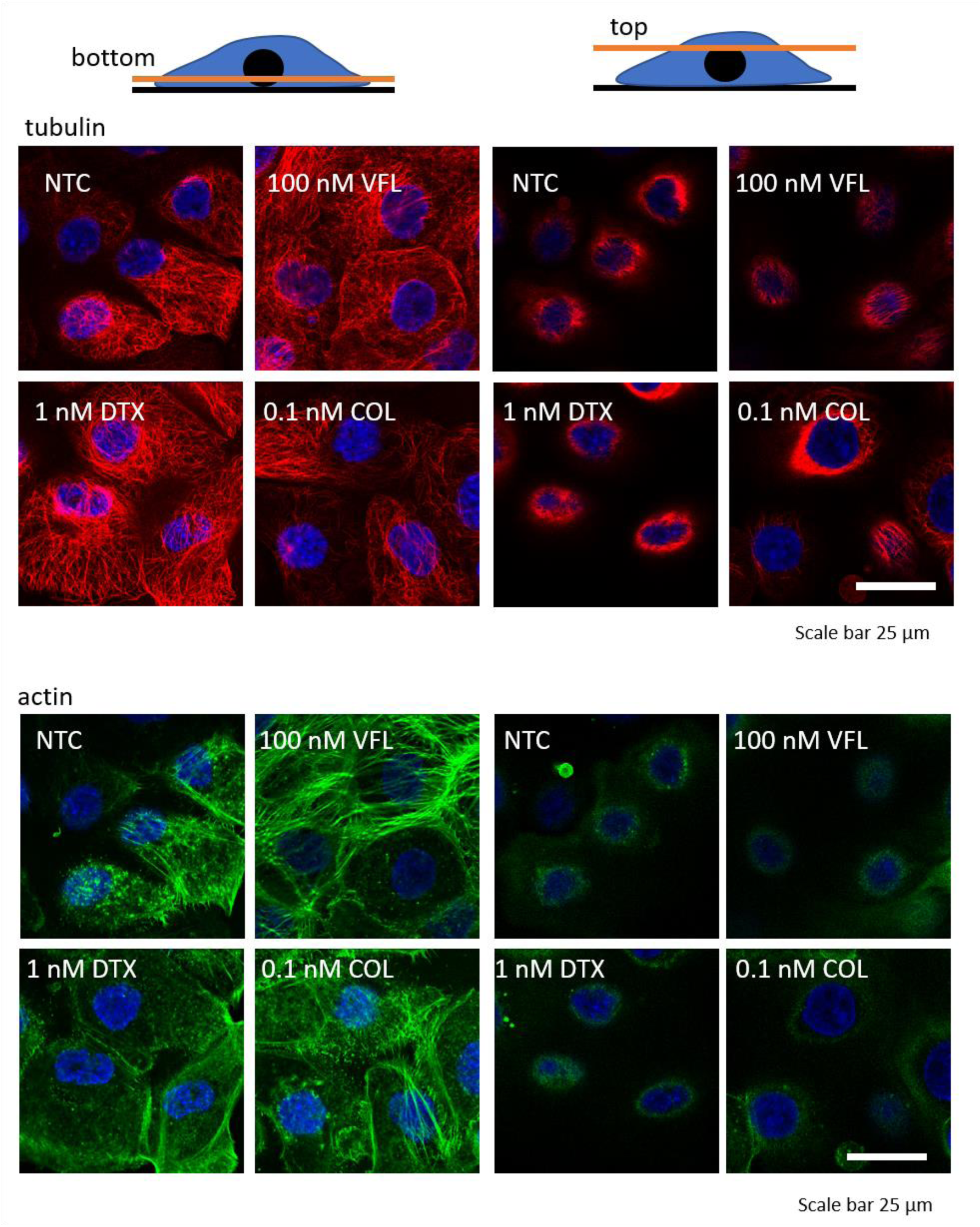
Confocal microscopy reveals a lack of cytoskeleton structural re-organization in microtubular networks linked with changes in mechanical properties of DU145 cells at low concentrations of MTDs. Fluorescent images were recorded at two positions along the cell height, top and bottom. The bottom position shows the organization of microtubules and actin filaments and microtubules in a focal plane close to the surface substrate. The top position shows the cytoskeleton organization at the top of the cell where the AFM probe touches its surface (F-actin - phalloidin-Alexa Fluor 488, microtubules β-tubulin antibody - Cy3, cell nuclei – Hoechst 33342).

## Discussion

Microtubules, and proteins associated with them, are critical for successful cell division, intracellular transport, signaling, and the regulation of focal adhesion dynamics both in normal and cancer cells. Their deregulation observed in cancer makes both tubulin and microtubules a prominent target for various chemotherapies^30^.

Numerous studies have reported a cell-, dose-, and time-dependent cellular response to microtubule-targeting drugs^6,31–35^. Although it is obvious that cells treated with high toxic concentrations are characterized by large damages causing their functional impairments, the effect of low drug concentrations on cells is less visible.

All MTDs interact with microtubules by binding to specific locations at the α/β tubulin heterodimer, thus, affecting the MTs dynamics by stabilizing or destabilizing the microtubular network^15,26,36,37^. Microtubule stabilizing and destabilizing drugs disrupt microtubule functioning, which is accomplished by various mechanisms.

Vinca alkaloids destabilize microtubules by binding to a site located at the inter-dimeric interface between *α* and *β* tubulin heterodimers^15^. They induce the formation of alternate microtubules leading to the dissociation of existing microtubules^38,39^. Taxanes reversibly bind to *β*-tubulin at the binding site located at the interior lumen of microtubules and enhance the polymerization process^40^. Colchicine binds to a site located at the interface between *α* and *β*-tubulin subunits, adjacent to the GTP-binding site of *α*-tubulin^41^. It blocks the availability of *α/β* tubulin heterodimers for protofilaments or MTs by changing the conformation of the dimers^32^. As a consequence, colchicine causes microtubule depolymerization by inhibiting lateral contacts between protofilaments^42^. Thus, we could expect that vinca alkaloids and colchicine should increase the softening of cells, while taxanes should enhance the stiffening of cells.

In this study, we have focused on the effect of three selected members of MTDs, i.e., DTX, VFL, and COL, on prostate DU145 cancer cells. Proliferation and MTS assays conducted for cells cultured on glass surfaces for low and high MTDs concentrations show confirmed dose-dependent effects. Applying high MTDs concentrations, regardless of their type, leads to cell death showing a significant decrease in the number of attached cells within a whole-time range (24h–72h). MTS assay carried out for cells substantially confirms the results of the proliferation assay. For low MTDs concentrations, a large drop in the number of attached cells visible after 72h of the exposure of cells to specific MTDs and weakly affected viability of cells indicated that the cell population is not uniform. It contains cells that detached from the surface and cells that still stay attached.

Calcein efflux demonstrated that VFL-treated cells are still able to pump out the drug, while this process is limited in the case of cells treated with DTX and COL (**Fig. 2e&f)**. These results indirectly show the accumulation of MTDs inside the cancer cells at drug concentrations below *IC*_*50*_. VFL in contrast to DTX and COL, impacts the cellular bioenergetics very weakly, which does not lead to drug accumulation. Disturbances within MTs and mitochondrial dynamics may be responsible for the diminution of intracellular energy deposits that are necessary for effective drug efflux by transmembrane protein complexes^43,44^.

The microtubular network alone has already been the object of AFM-based nanomechanical studies. In most cases, research was focused on the mechanical properties of cells treated by taxol-based compounds^19,20,45–47^. Less frequently, the effect of compounds such as COL or vinca alkaloids, has been evaluated using AFM^46,48^. Results gathered from a few existing publications report no stiffness change for various cell types treated with COL within a time frame of 2 h and 4 h for applied concentrations of 500 µM and 10 µM, respectively^46,48^. Vincristine (belonging to vinca alkaloids) induced stiffening of cells treated with 3 nM, 12 nM and 60 nM of drug concentrations for 48 hours^47^. Paclitaxel (belonging to taxanes) affects mechanical properties of cells in various manners – in some cells no effect was observed^46^ while in others an increasing cell rigidity was detected^21,47,48^. Compared to these results, our findings are different, which might be due to a distinct response of prostate DU145 cells to MTDs as other studies have already demonstrated the susceptibility of cancer cells to cell death induced by these drugs^33,34^.

According to our results, low concentrations of MTDs applied to prostate DU145 cancer cells induced changes in the organization of the microtubular network and the mechanical properties of cells in a surprising manner. For the VFL, which weakly affected the MTs organization, the largest cell stiffening is observed. Cells became more rigid. DTX-treated cells soften at high (above *IC*_*50*_) concentrations at which large damages of MTs are detected. For COL, which had the strongest effect on the microtubular network, the mechanical properties of cells remained almost unchanged.

From these data, we cannot support the hypothesis that the mechanical resistance of prostate cells is associated with the changes in the organization of microtubules. Nor from the impairments in efflux pumps that transport the drugs out of the cancer cells^43,44^. We postulate that the stiffening of prostate DU145 cancer cells is likely the mechanism that helps cancer cells to function in a non-disturbed way (like there were no MTDs presence).

Looking at the time-related change of cell deformability (**Fig. 4a-c)**, all MTDs induce the maximum deformability changes in cells within a defined time of exposure to them. By analyzing MTDs-induced changes in both actin and microtubular networks, we noticed the formation of actin bundles in the affected cells. In our data, stiffening in VFL-treated cells was accompanied by the enhanced formation of actin bundles (**Fig. 6; Suppl. Fig. S5**). DU145 cells treated with DTX and COL for the same time reveal the mechanical properties of the level of NTC. For these cells, no increased number of actin bundles was observed indicating that the crosstalk between actin and microtubular network was significantly weaker. Our findings show the crosstalk between actin filaments and microtubules is likely the mechanisms that could be involved in the resistance of cells to vinca alkaloids observed in chemotherapy.

## Materials and Methods

### Microtubule-targeting drugs (MTDs)

The MTDs chosen for our study were vinflunine, docetaxel, and colchicine. Vinflunine ditartrate (VFL, Toronto Research Chemicals) was dissolved in deionized water (Cobrabid purification system, 18 Ωcm^-1^) to prepare stock solutions of 1 mM vinflunine concentration. They were stored in -18°C. Docetaxel and colchicine (DTX and COL, respectively, Sigma-Aldrich) were dissolved in ethanol. Drugs were added to the cell cultures 24 hours after cell seeding, followed by the application of a set of methods (**Fig. 1a)** to characterize properties of prostate cancer cells.

### Cell culture

The human prostate cancer cells (DU145, ATCC, LGC Standards) were cultured in Dulbecco’s Modified Eagle’s Medium (DMEM, Sigma-Aldrich) supplemented with 10% heat-inactivated Fetal Bovine Serum (FBS, LGC Standards) without antibiotics. Cells were grown in 25 cm^2^ culture flasks (Saarstedt), passaged every 3-4 days by trypsinization with trypsin-EDTA solution (Sigma-Aldrich) and moved to the corresponding Petri dishes. To study the effect of anticancer drugs targeting microtubules, VFL, DTX, and COL were added to cells cultured in Petri dishes. Final concentrations of anticancer drugs applied to cells were 100 nM and 750 nM for VFL, 1 nM and 5 nM for DTX, 0.1 nM and 0.5 nM for COL. As a control, non-treated cells were used.

### Assessment of drugs cytotoxicity by MTS assay

Cytotoxicity of drugs was assessed by Cell Growth Determination Kit, an MTT based colorimetric assay (Sigma-Aldrich). 3000 of DU145 cells/per well were seeded on 96-well cell culture plate (TPP). After the overnight culture of cells, the culture medium was exchanged to that containing the corresponding drug (VFL, DTX, or COL). After 68 hours of culture with drugs, 10 µl of 3-(4,5-dimethylthiazol-2-yl)-2,5-diphenyltetrazolium bromide was added to each well. After 4 hours of incubation (at time point 72h), formazan crystals were dissolved in DMSO and absorbance was measured by spectrophotometer (ELISA SPECTROstar Nano, BMG LABTECH) at 570 nm wavelength for dissolved formazan and 690 nm for a background.

### Proliferation of cells

To assess the effect of anticancer drugs on DU145 cell proliferation, cells were seeded at the density of 3000 cells/cm^2^ in cell culture dishes with Ø = 35 mm (Sarstedt). They were kept in DMEM medium containing no phenol red (Sigma-Aldrich), 10% heat-inactivated FBS, and 2 mM L-glutamate (Sigma-Aldrich). The same number of cells was applied, i.e. 3 000 cells per ml for all samples. After 24 hours, the culture medium was replaced by a medium containing a specific drug concentration. Then, cells were maintained in such a medium for 24 h, 48 h, and 72 h. At each time point, a part of the cells was passaged and counted using the Bürker chamber.

### Atomic force microscopy – indentation mode

For the measurements of the relative changes of the mechanical properties of the cells at different time-points, AFM indentation experiments were carried out using both pyramidal and spherical probes. It has been already shown that indentation data of living cells collected with sharp and spherical probes can both detect changes between two cell populations^49^. Mechanical properties of DU145 cells were determined by analyzing force curves (additional experimental details are included in **Suppl. Notes S3 and S4**). The raw force curves (raw cantilever deflection versus z-piezo displacement) were converted into force versus indentation curves in the proper units (nN and nm, respectively) according to consolidated procedures^50^. The spring constant of the cantilever was measured using the thermal noise calibration^51^. The deflection sensitivity was calibrated by recording raw force curves on non-deformable substrates (coverslip without cells in this case) and evaluating the inverse of the slope of the uncalibrated contact region^50^. Alternatively, the deflection sensitivity was also calibrated in situ and non-invasively by imposing that the result of the thermal noise calibrated should be compatible with the previously characterized spring constant, according to the SNAP procedure^52^. Assuming that the elastic response of the cell can be described by Hertz contact mechanics^53^, Young’s modulus was determined.

In the case of a conical approximation of the probing tip, the relation between load force *F* and the resulting indentation *δ* is:

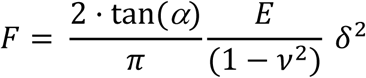

where *α* is the open-angle of the probing cone, *ν* is the Poisson’s ratio, and *E* is the effective Young’s modulus. For a sphere indenting purely elastic material, the relation between load force *F* and the resulting indentation *δ* is:

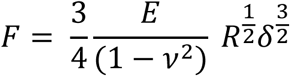

Where *R* is the probe’s radius. All calculations were carried out assuming that *ν* = 0.5 (incompressible cells).

### Characterizing mechanical properties of different cellular regions

To evaluate Young’s modulus changes occurring at different regions of the cell, information on both a local height of the cell body and the value of the apparent Young’s modulus was determined from each single force curve, and further used to provide the topographical and the Young’s modulus maps^23^ (see **Suppl. Note S4**). A finite-thickness correction was applied to extract accurate values of the Young’s modulus from the force curves^54^ coming from experiments with colloidal probes because the rigid substrate underneath the cells can cause an apparent increase of the Young’s modulus that depends on the local height of the sample, therefore masking fine mechanical modifications that take place inside the cells body. The correction was not applied to pyramidal sharp tips since the finite-thickness effect becomes relevant with big contact area (see **Suppl. Note S5**).

Force curves were linearized to identify the presence of multiple elastic regimes inside the cell (typically attributed to an upper layer made of the actin cortex, and to an inner layer including the region around the cell nucleus). Two linear regions were typically identified in the linearized curves: a first region corresponding to the 5-15% relative indentation range (to the local values of cells height), the shallow indentation range, and a second inner region, corresponding to the 20-40% relative indentation range, the deep indentation range. Then, the Hertz model with applied finite thickness corrections was applied to the force curves in the two indentation ranges, obtaining two maps, representing the rigidity of the surface and inner part of the cell, respectively (**Fig. 5, Suppl. Table 2)**.

To characterize the elastic properties of different parts of the cell’s body we created masks to select force curves belonging only to specific regions of interest (nuclear and peripheral region with varying indentation). Nuclei are usually located in the tallest cell region. Since the nucleus elasticity could be hidden by that of the stiff actin cortex, we decided to build the mask by multiplying point by point the topographic and the elasticity maps. We found this procedure more reliable than identifying the nuclei simply as belonging to the tallest regions in the topographic maps. The masks allowing to segregate the force curves belonging to the nuclear regions were built exploiting both the elastic and the topographical maps **as shown in Fig. 5**.

### Statistical analysis for the determination of Young’s modulus for given cellular regions

The Young’s modulus of each cellular region typically follows a lognormal distribution, which appears as Gaussian distributions in semilog10 scale. A Gaussian fit in semilog10 scale provides the peak value *E*’ and the geometric standard deviation *σ*_*g*_^*10*^, from which the median *E*_*med*_ and the standard deviation of the median *σ*_*med*_ can be calculated ^59^:

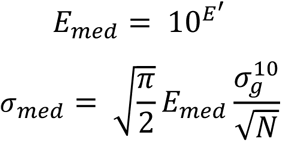

where *N* is the number of force curves recorded for each cell cluster. The total error for the modulus median *σ*_*clust*_ associated to each cluster was calculated by summing in quadrature *σ*_*med*_ and a calculated instrumental error *σ*_*inst*_ = 3%^37^:

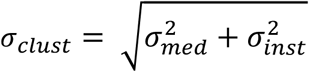

Then, the median mean *E*_*mean*_ of the moduli median, measured for different clusters at a given condition for the different cellular regions, was evaluated as: 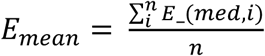 where *i* is the number of clusters investigated. The total error *σ*_*E*_ associated to *E*_*mean*_, representative for a specific condition and cellular region, was calculated by summing in quadrature the propagated error of the medians *σ*_*S*_ and the standard deviation of the mean *σ*_*mean*_:

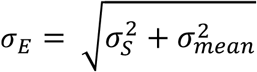

where

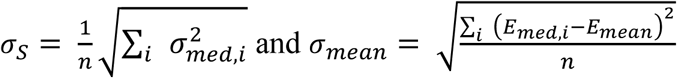

and *n* is the number of clusters investigated.

### Immunofluorescence staining

To assess the impact of microtubule-targeting drugs on the organization of the microtubule network in DU145 cells fluorescent microscopy was employed. Cells were seeded in density 3000 cells/cm^2^ on glass coverslips and allowed to stabilize at substrate overnight. Afterward, the culture medium was exchanged to a fresh medium containing final concentrations of microtubule-targeting drugs. After 72 hours, the culture medium was removed and cells were rinsed 3 times with Dulbecco’s Phosphate Buffered Saline (DPBS, Sigma-Aldrich).

Next, cells were fixed with 3.7% formaldehyde solution and washed with DPBS to remove remaining formaldehyde. Permeabilization of cells membranes was realized by the treatment of fixed cells with 0.2% Triton X-100 solution, followed by washing with DPBS. Actin filaments were stained with phalloidin conjugated with Alexa Fluor 488 dye (0.33 μM) for 45 minutes. Afterward, cells were incubated overnight with anti-β-tubulin antibody conjugated with fluorescent dye Cy3 (1:200) (Sigma-Aldrich), followed by nuclei counterstaining using Hoechst 33342 (0.2 μg/mL).

### Fluorescent and confocal microscopy

Epi-fluorescent images were acquired using a fluorescent microscope (Olympus IX53) equipped with fluorescence camera XC10 (1.4 mln pixels). The focal plane was set close to the cell surface. High-resolution 2D and 3D images recorded at the focal planes close to the glass surface and in the apical sections (where AFM probed mechanical properties of cells) were acquired by Leica TCS SP8 X confocal laser-scanning microscope (Leica Microsystems CMS GmbH, Germany) using 63x HC PL APO CS2 1.40 NA oil immersion objective. Image acquisition was performed bidirectionally along X-axis at 200 Hz scan speed and image format 1024×1024 (pixel size 72 nm), zoom 2.5, and line average 2. Three-dimensional images were reconstructed using Leica Application Suite X software (3D Movie Editor, Leica Microsystems CMS GmbH, Germany) or ImageJ 1.51n (Wayne Rasband, National Institute of Health, USA) on default parameters (except non-linear intensity adjustment – gamma parameter was 0.6).

### Calcein efflux monitoring

Cells were seeded into 6-well plates at a density of 60 000 cells/cm^2^ and treated with proper concentrations of MTDs for 24 h. After incubation with 0.75 µM calcein-AM (Sigma-Aldrich) for 30 min, the cells were washed with a warm PBS buffer and incubated in a fresh medium for another 3h. After harvesting with trypsin/EDTA solution, the cells were centrifuged and re-suspended in FluoroBrite® DMEM (supplemented with 10% FBS and 1% GlutaMAX; Gibco) and acquired with ImageStreamX MkII system and ISX software (Luminex Corp.) to determine the efficiency of dye efflux. Data processing was performed with IDEAS 6.2 software. The analysis was done on “*in-focus*” images of single cells. In-focus single cells were classified based on their bright-field area and high bright-field aspect ratio (width to height ratio). At least 10 000 images were collected for data analysis.

### Statistics

For cell biology methods (proliferation and MTS assay, cytometry, morphometric analysis) data are presented as a mean ± standard deviation (s.d) (**Fig. 1b-e, Fig. 2e-f**) calculated from three independent experiments. Data describing the morphometric properties of cells (**Suppl. Fig. S1 and Suppl. Fig. S2**) are expressed as a mean ± s.d. calculated for *n* denoting the number of cells measured (specified in the figure legends). Original Young’s modulus values were calculated as a mean ± s.d considering the number of cells or clusters measured (**Suppl. Tables S1-2)**. Relative Young’s modulus changes presented in **Figs. 4, 5, Suppl. Figs. 2-3)** were accompanied by an error calculated using the error propagation method. In all cases, statistical significance was estimated by unpaired t-Student test at the level of 0.05. The following notation was applied: * p < 0.5; ** p < 0.1; *** p < 0.001.

### Data availability

The authors declare that the data supporting the findings of this study are available within the paper and its supplementary information file. Raw data underlying the presented figures are available from the corresponding authors upon reasonable request.

## Acknowledgments

AK and NB acknowledge the support of InterDokMed project no. POWR.03.02.00-00-I013/16. AP and ML acknowledge the support of the European Union under the Marie Skłodowska-Curie grant agreement No. 812772 (Phys2BioMed). AP and CS acknowledge the support of the European Union under the FET Open grant agreement No. 801126 (EDIT). DR and MP acknowledge the support of the Polish National Science Centre (grant no. UMO-2015/19/D/NZ3/00273). ML & AK are grateful to Mrs. Klaudia Suchy for help in cell cultures and to Dr. Justyna Bobrowska in help in recording epi-fluorescence imaging of cell nucleus.

