## Supplementary Materials for "Stiffening of prostate cancer cells driven by actin filaments – microtubules crosstalk confers resistance to microtubule-targeting drugs"

**Content of the supplementary information:**


- ***Supplementary Note 1: Proliferation rate of DU145 cells treated with MTDs***

- ***Supplementary Note 2: Morphology of DU145 cell nucleus is affected by MTDs (****Supplementary Figure S1****;*** *Supplementary Figure S1)*

- ***Supplementary Note 3: Time-dependent changes in mechanical properties of DU145 cells (****Supplementary Table S1; Supplementary Figure S3)*

- ***Supplementary Note 4: Mapping the mechanical properties of cells***

- ***Supplementary Note 5. Finite-thickness correction*** *(Supplementary Figure S4)*

- ***Supplementary Note 6: 3D organization of MTs visualized using confocal microscopy***

- ***Supplementary Note 7: Actin organization inside DU145 cells (****Supplementary Figure S5, Supplementary Figure S6)*

***Supplementary Note 1: Proliferation rate of DU145 cells treated with MTDs***

To estimate how fast the number of cells increases, a linear regression was applied to quantify roughly the rate of proliferation **(Fig. 1)**. Although for NTC cells an exponential-like growth can be observed, the linear fit was chosen to avoid an exponential fit to three points. As expected, NTC cells had the highest proliferation rate, which was of 4300 ± 1100 cells per hour (Pearson's' coefficient = 0.939). For low doses of drugs, i.e. 100 nM VFL, 1 nM DTX and 0.1 nM COL, obtained rates were 3000 ± 270 cells per hours (Pearson's' coefficient = 0.992), 2270 ± 220 cells per hour (Pearson's' coefficient = 0.991) and 3340 ± 740 cells per hour (Pearson's' coefficient 0.954), respectively.

***Supplementary Note 2: Morphology of DU145 cell nucleus is affected by MTDs***

To investigate nuclear morphology fluorescence microscopic images were analyzed in ImageJ with package Fiji and Cookbook with the function Particle Analysis. At least 5 images for each group for the following drugs concentrations: 100 nM and 750 nM for VFL, 1 nM and 5 nM for DTX, 0.1 nM and 0.5 nM for COL and control without addition of drug.

Although observed changes in microtubule organization correspond roughly with the results of cell viability, we wanted to verify the morphology of the cell nucleus (**Suppl. Fig. S1**). At large drug doses, regardless of the drug applied, cell nuclei were heavily irregular and defragmented as one can see in the Hoechst stained fluorescent images (**Suppl. Fig. S1a**).

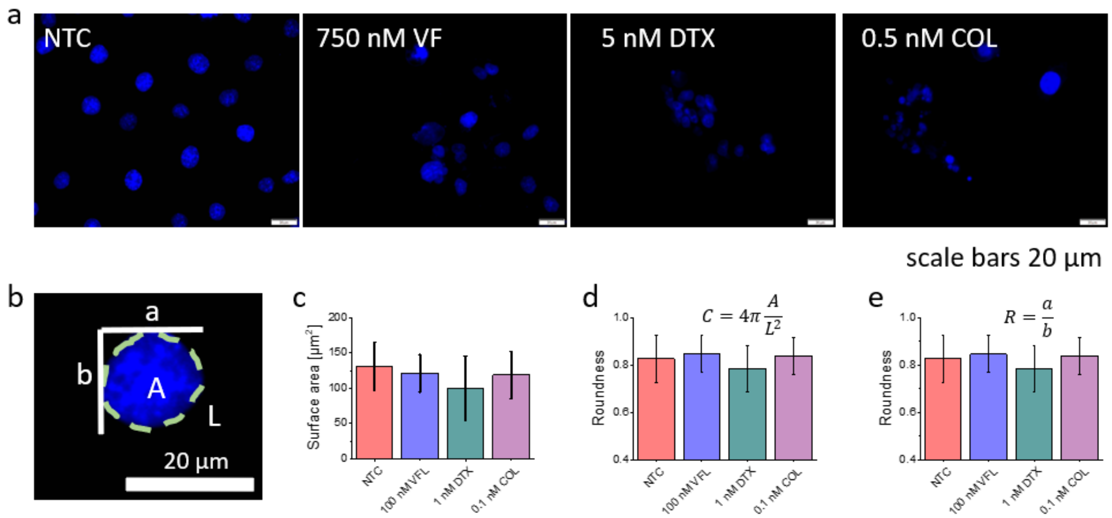


***Suppl. Figure S1***. *Morphology of the cell nucleus is strongly affected by microtubule-targeted drugs.* ***a***, *High concentrations of VFL, DTX, and CL induce fragmentation of cell nuclei, especially for DTX and COL. In low drug concentration, nucleus morphology is preserved (****b****), however, DTX seems to affect nucleus changes, as shown in morphometric analysis of surface area (****c****), circularity (****d****), and roundness (****e****) for untreated and treated prostate DU145 cells. Data are presented as a mean ± standard deviation calculated from n = 79-105 cells per condition.*

To evaluate nuclear changes occurring at low doses, three morphometric parameters were calculated, namely, nuclear surface area, circularity, and roundness (**Suppl. Fig. S2**). Obtained results showed large variations in the nucleus shape, in particular for DTX treated cells, for which circularity and roundness values dropped, too. For cells treated with COL and VFL cell nuclei seem to be less affected by the MTDs action.

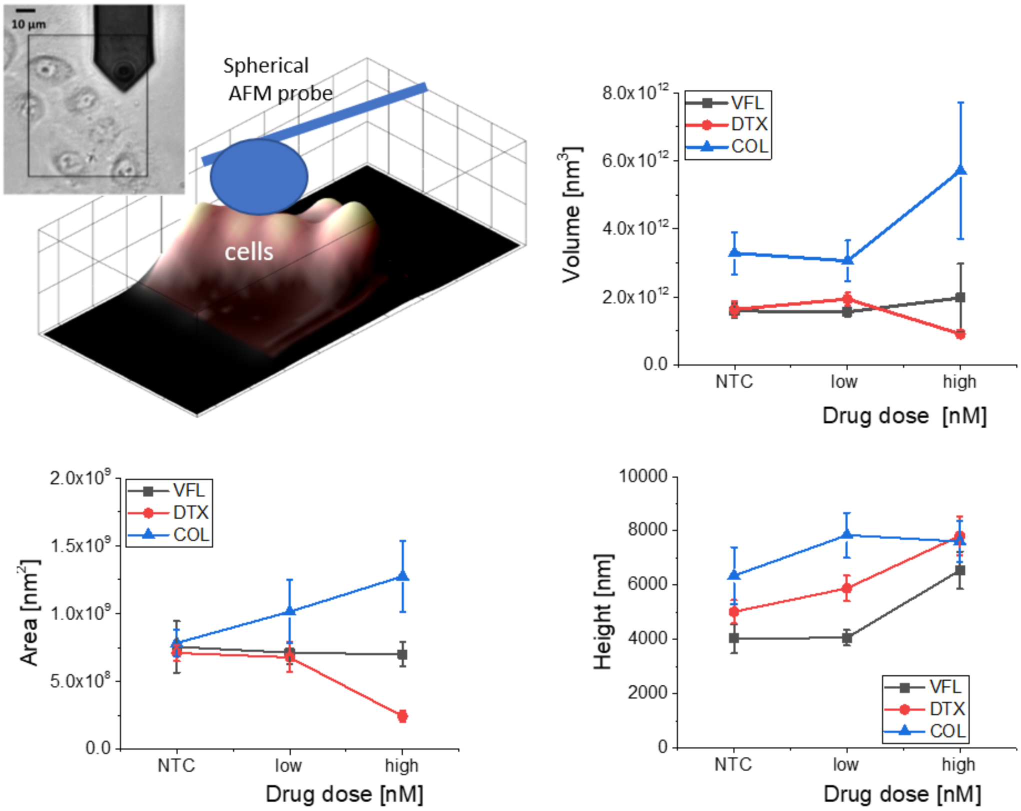


***Suppl. Figure S2. Cells change their morphology in response to MTDs.*** *AFM-based morphometric parameters of DU145 clusters showing alterations in clusters volume, surface area, and height induced by MTDs. Data are presented as a mean ± $\sigma_E$; n = 8-12 per condition.*

The obtained results show significant and opposing alterations of the cell volume and surface area for DTX (decrease) and COL (increase), especially for the high doses. For high VFL concentration, a significant increase in total volume in comparison to NTC is also observed. Such observation is leading us to the conclusion that the swelling of cells treated with a high concentration of drugs is characteristic of microtubule-disrupting agents. The height of the analyzed cell clusters shows an increasing trend for all MTDs, although significant observations were unique for high drug concentrations. Interestingly, in the low-dose (100 nM) VFL case, for which pronounced deformability changes were recorded, volume, surface area, and height of cells remained at the same level of the NTC cells. Thus indicating cytoskeleton contribution in mechanical features changes of 100 nM VFL treated cells, rather than changes in their dimensions.

***Supplementary Note 3: Time-dependent changes in mechanical properties of DU145 cells***

To measure the mechanical properties of DU145 cells, cells were seeded on glass coverslips (Ø = 25 mm) immersed in a Petri dish with internal Ø = 34 mm (TPP) at the density of 50 000/ml. Measurements were carried out using atomic force microscopes: model Xe120 (Park Systems) and working in a force spectroscopy mode. A rectangular cantilever ORC8 (Bruker) with a nominal spring constant of 0.05 N/m, the opening angle of 36 ± 2º, and a radius of 15 nm were used. Cantilever spring constants were calibrated by the use of the thermal noise method[1]. For measurements, coverslips with cells were mounted in a transparent liquid cell and kept at room temperature in standard culture medium. Room temperature was not significantly harmful for Du145 during experiments which lasted up to 2 hours[2]. Keeping a temperature below 37°C reduces the cell motility, which was beneficial for the elasticity measurements. 25 force curves were acquired within an elasticity map of 5 × 5 points with a pixel size of 1 $\mu m^2$, load force ~10 nN, and tip velocity of 8 µm/s. Each experiment was conducted in triplicate.

The relative Young's modulus variations are, calculated as $\Delta E_{rel} = (E_{drug}-E_{NTC})/E_{NTC}$, are reported in **Suppl. Table S1 and Suppl. Fig. S2.** In our analysis, the maximum indentation was 500 nm. The final Young's modulus was calculated as the mean ± standard error (s.e.) from *n* = 80-96 cells measured for each group separately (**Suppl. Table S1)**.

*Suppl. Table S1. Young's modulus obtained for MTDs treated Du145 cells (s.e. – standard error, n – number of cells measured; p-value was calculated using two tails unpair Student t-test). The relative Young's modulus variations are reported in both Fig. 4 and Suppl. Fig. S2.*

| ***time*** / ***MTDs treatment*** | ***24h***<br>*E ± s.e. (n) [kPa]* | ***48h***<br>*E ± s.e. (n) [kPa]* | ***72h***<br>*E ± s.e. (n) [kPa]* |
|---|---|---|---|
| ***Vinflunine (VFL)*** | | | |
| ***NTC*** | 3.1 ± 0.2 (94) | 2.2 ± 0.1 (93) | 1.5 ± 0.1 (96) |
| ***100 nM VFL*** | 3.1 ± 0.3 (94)<br>*p = 0.9512* | 3.3 ± 0.3 (93)<br>*p = 0.0001* | 1.9 ± 0.1 (93)<br>*p = 0.0016* |
| ***750 nM VFL*** | 3.0 ± 0.2 (76)<br>*p = 0.7147* | 2.7 ± 0.2 (88)<br>*p = 0.0412* | 2.5 ± 0.2 (91)<br>*p = 0.0001* |
| ***Docetaxel (DTX)*** | | | |
| ***NTC*** | 4.2 ± 0.3 (88) | 2.9 ± 0.2 (88) | 2.6 ± 0.1 (89) |
| ***1 nM DTX*** | 4.7 ± 0.4 (86)<br>*p = 0.3610* | 4.8 ± 0.4 (83)<br>*p = 0.0001* | 2.8 ± 0.2 (89)<br>*p = 2977* |
| ***5 nM DTX*** | 4.4 ± 0.4 (83)<br>*p = 0.7280* | 3.4 ± 0.3 (86)<br>*p = 0.1085* | 2.8 ± 0.1 (87)<br>*p = 0.2971* |
| ***Colchicine (COL)*** | | | |
| ***NTC*** | 3.3 ± 0.17 (90) | 3.0 ± 0.15 (91) | 2.2 ± 0.1 (90) |
| ***0.1 nM COL*** | 4.4 ± 0.3 (88)<br>*p = 0.0012* | 2.9 ± 0.2 (86)<br>*p = 0.7853* | 2.3 ± 0.1 (87)<br>*p = 0.4623* |
| ***0.5 nM COL*** | 3.3 ± 0.2 (87)<br>*p = 0.9140* | 2.0 ± 0.1 (88)<br>*p = 0.0001* | 2.0 ± 0.2 (88)<br>*p = 0.2520* |

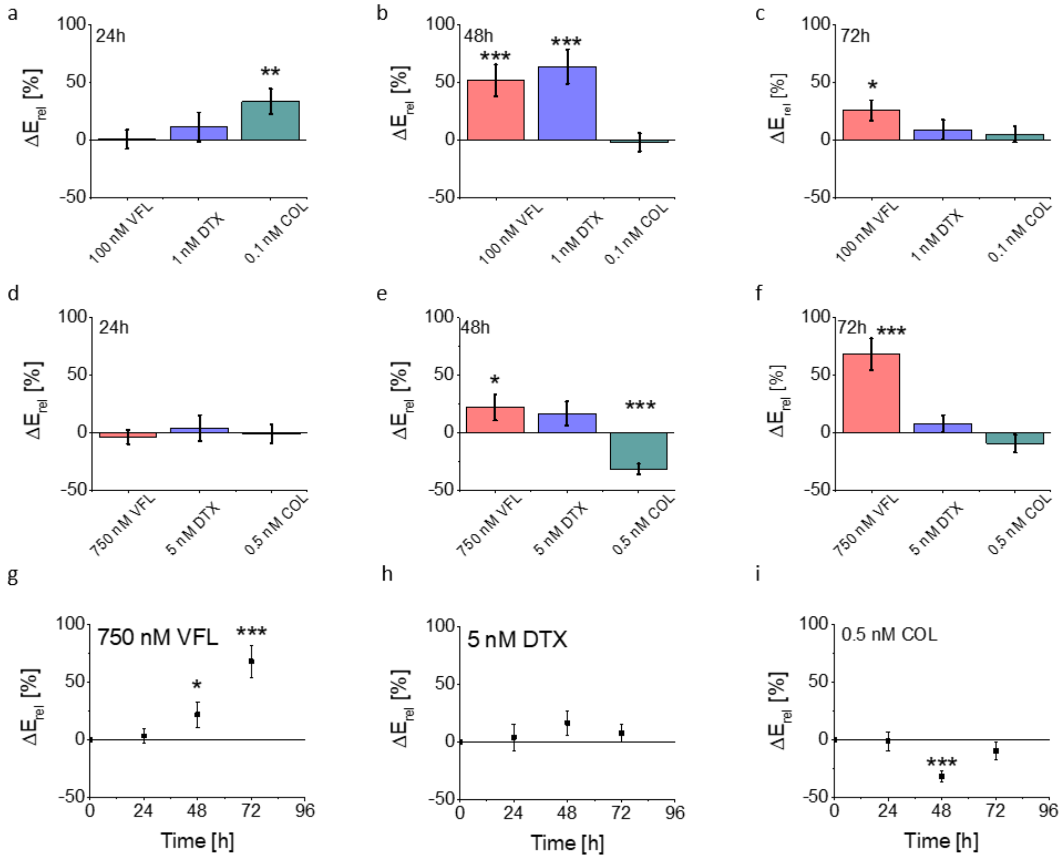


***Suppl. Figure S3**. Relative change $\Delta E_{rel}$ of the Young's modulus measured at the same time-point for low (**a-c**) and high **(d-f)** MTDs doses. **(g-i)** Time-dependent $\Delta E_{rel}$ changes showing cell stiffening or softening dependently of the MT drug, concentration, and time applied. Error bars were calculated using the error propagation method. Statistical significance was estimated by unpaired t-Student test at the level of 0.05 (* $p < 0.05$; ** $p < 0.01$).*

### *Supplementary Note 4: Mapping the mechanical properties of cells: combined topographical and mechanical mapping*

For acquiring maps of mechanical properties, DU145 cells were plated in the standard experimental conditions on Ø = 40 mm glass-bottom dishes for cell culture (Willco Wells, Amsterdam, Netherlands). Cells were maintained in DMEM without phenol red but supplemented with 10% heat-inactivated FBS and 5 nM L-glutamate. Glass coverslips with cells were mounted in transparent liquid cell and kept at room temperature in standard culture medium.

Combined topographical and mechanical imaging was performed using a Bioscope Catalyst AFM (Bruker). During the AFM measurements, the temperature of the medium was maintained at 30°C using a perfusion stage incubator and a temperature controller (Lakeshore 3301, Ohio, USA). The choice of the temperature below 37°C was made to decrease the cellular metabolism, which slows down also the cell motility, making it possible to acquire large maps without disturbance due to strong movements of the cells during the measurements (acquisition of one map takes 30-40 min).

We used custom monolithic borosilicate glass probes consisting in spherical glass beads with radii $R$ in the range 4500–5500 nm, attached to ultra-soft silicon, tipless cantilevers (Micromash HQ:CSC38/tipless/no Al) with elastic constant k = 0.02– 0.03 N/m, measured using the thermal noise calibration[3]. Probes were fabricated and calibrated, in terms of tip radius, according to an established custom protocol[4].

The topographical/mechanical maps were obtained, as described elsewhere[5], from sets of force curves, collected in Point and Shoot (P&S) mode, selecting the regions of interest from optical images, exploiting the accurate alignment of the optical and AFM images obtained using the Miro software module integrated in the AFM software. Each set of force curves consisted of an $N \times M$ array of curves spatially separated by approximately 1.5 µm, each force curve containing 4096 points, with ramp length $L$ = 6 µm, maximum load $F_{max}$ = 10 nN, ramp frequency $f$ = 1 Hz. Typical maximum indentation was 2-3 µm, larger than in the case of sharp pyramidal probes (max indentation 500 nm).

*Suppl. Table S2. Young's modulus obtained for MTDs treated DU145 cells showing the mean value of the median E extracted per each cluster; error represents the final error $\sigma_E$ that take into account the three main errors: the instrumental error $\sigma_{inst}$, the error associated to one single cluster median value $\sigma_{med}$ and the deviation of the all clusters $\sigma_{mean}$, n – number of cell clusters measured; p-value was calculated using the two-tails unpair Student t-test.*

| | ***VFL*** | ***DTX*** | ***COL*** |
|---|---|---|---|
| ***whole-cell*** | *E ± $\sigma_E$ [Pa]* | *E ± $\sigma_E$ [Pa]* | *E ± $\sigma_E$ [Pa]* |
| ***NTC*** | 375.1 ± 23.2 (8) | 429.0 ± 35.9 (11) | 376.6 ± 37.6 (9) |
| ***low*** | 444.61 ± 24.4 (8)<br>*p = 0.0058* | 407.9 ± 34.3 (12)<br>*p = 0.6759* | 360.6 ± 79.6 (8)<br>*p = 0.8528* |
| ***high*** | 447.1 ± 29.8 (7)<br>*p = 0.0755* | 336.6 ± 25.5 (12)<br>*p = 0.0456* | 436.8 ± 132.1 (7)<br>*p = 0.6321* |
| ***nuclear region (5-15%)*** | | | |
| ***NTC*** | 346.6 ± 19.8 (8) | 410.8 ± 32.7 (11) | 422.9 ± 43.6 (9) |
| ***low*** | 466.5 ± 36.9 (8)<br>*p = 0.0125* | 388.3 ± 41.0 (12)<br>*p = 0.6760* | 470.5 ± 99.2 (8)<br>*p = 0.6536* |
| ***high*** | 539.0 ± 44.9 (7)<br>*p = 0.0012* | 394.4 ± 33.6 (12)<br>*p = 0.7296* | 628.4 ± 167.9 (7)<br>*p = 0.2054* |
| ***nuclear region (20-40%)*** | | | |
| ***NTC*** | 381.9 ± 13.2 (8) | 378.7 ± 24.3 (11) | 429.0 ± 38.2 (9) |
| ***low*** | 488.9 ± 22.7 (8)<br>*p = 0.011* | 341.8 ± 25.5 (12)<br>*p = 0.3093* | 486.1 ± 60.6 (8)<br>*p = 0.4271* |
| ***high*** | 537.6 ± 44.8 (7)<br>*p = 0.0037* | 340.0 ± 16.5 (12)<br>*p = 0.1957* | 642 ± 107 (7)<br>*p = 0.0584* |
| ***cell periphery*** | | | |
| ***NTC*** | 397.3 ± 29.9 (8) | 401.4 ± 27.6 (11) | 376.8 ± 35.9 (9) |
| ***low*** | 437.5 ± 24.2 (8)<br>*p = 0.3139* | 403.4 ± 23.1 (12)<br>*p = 0.9544* | 388.5 ± 79.3 (8)<br>*p = 0.8905* |
| ***high*** | 398.3 ± 41.7 (7)<br>*p = 0.9841* | 321.1 ± 24.8 (12)<br>*p = 0.0416* | 536 ± 152 (7)<br>*p = 0.2702* |

***Supplementary Note 5. Finite-thickness correction***

To overcome issues linked with the influence of the underlying stiff support on the mechanical properties of cells, finite thickness correction was applied to the obtained data shown in **Suppl. Fig. S4**.

The finite-thickness correction is needed when the requirement that the sample thickness is much larger than the maximum indentation ($h >> \delta$) is not satisfied. When indenting cells, which are very soft, with maximum indentation $\delta$ well beyond 1 µm, the finite-thickness effect must be accounted for. In this condition, the influence of the rigid substrate (the bottom of the glass dish from Wilco Weels®) makes the cell elastic response stiffer, i.e. the measured YM is larger. This effect increases when the sample becomes even thinner, like in the peripheral regions of the cell body. As a rule of thumb, the condition 10-15 $\delta < h$ should be satisfied to neglect finite-thickness effects, although the incidence of finite thickness depends also on the radius of the indenter (see below). It turns out actually that this effect is stronger for large colloidal probes, while it is usually negligible for sharp AFM tips.

We implemented the analytic correction developed by Dimitriadis *et al.*[6] for spherical probes, as described by Puricelli *et al.*[5]. Since cells adhere to the substrates using highly dynamical focal adhesion complexes, following the advice of Gavara and Chadwick[7], we used a correction factor to the Hertz equation, which represents an average between the two conditions of perfectly adherent and non-adherent cells:

$$F = \frac{9}{16} E R^{\frac{1}{2}} \delta^{\frac{3}{2}} [1 + 1.009\chi + 1.032\chi^2 + 0.578\chi^3 + 0.051\chi^4]$$

This is the effective equation for the correction of the finite-thickness effect.
Defining $\Delta(\chi) \equiv \Delta(\chi(R, \delta, h))$, with

$$\Delta = 1 + 1.009\chi + 1.032\chi^2 + 0.578\chi^3 + 0.051\chi^4,$$

the finite-thickness corrected Hertz Model can be written as:

$$\frac{F}{\Delta(\chi)} = \frac{9}{16} E R^{\frac{1}{2}} \delta^{\frac{3}{2}}$$

The variable $\chi = \frac{\sqrt{R\delta}}{h}$ combines the three critical lengths of the system, the radius of the probe R, the indentation $\delta$ and the height of the sample $h$. Considering that for the Hertz model the contact area between the probe and the sample is $a = \sqrt{R\delta}$, we have $\chi = \frac{a}{h}$, meaning that the finite-thickness effect does not depend directly on the ratio of the vertical lengths $\delta$ and $h$, but rather on the ratio of the horizontal dimension of the contact area, i.e. the contact radius $a$, to the sample height $h$. As a consequence, for a given indentation δ, the finite-thickness effects will be significantly stronger for the colloidal probes, compared to the sharp ones.

An example of the effects of the finite-thickness correction is shown in **Suppl. Figure S4**. The value of the Young's modulus extracted from each force curve is plotted as a function of the local cell height, for three representative different clusters of cells (control conditions), measured in a single day. Data are shown with (**Suppl. Fig. S4a)**, or without (**Suppl. Fig. S4b)**, the finite-thickness correction.

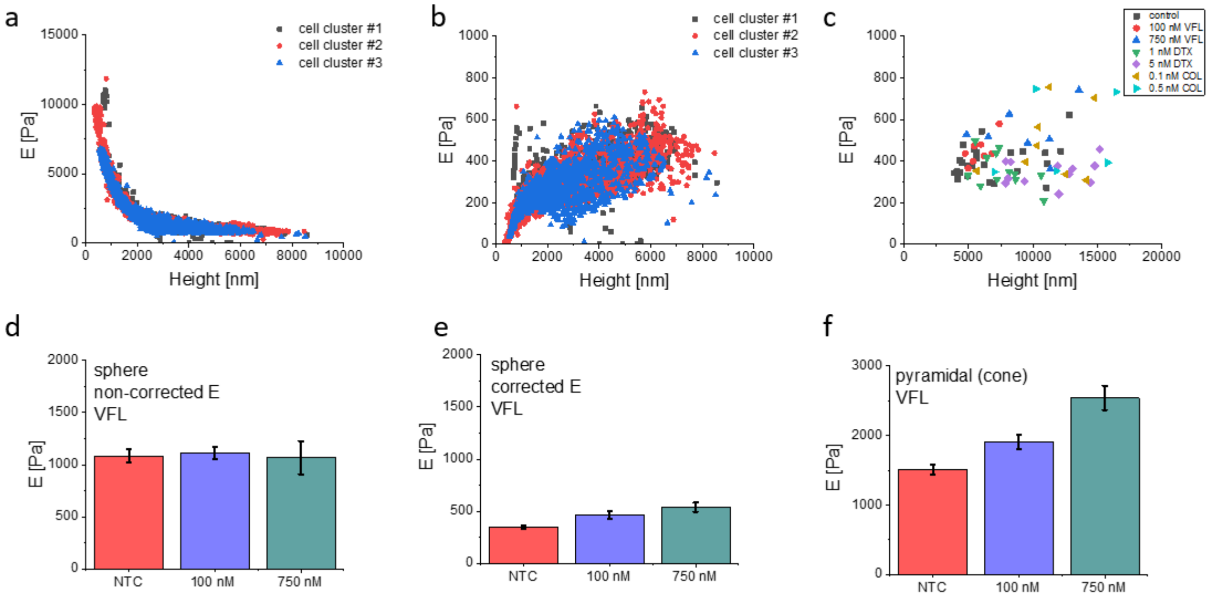


***Suppl. Figure S4***. *Finite thickness correction for colloidal probes based on Dimitriadis et al.*[6] *Data are presented as Young's modulus extracted from single force curves, coming from three clusters of cells of the control condition, using (a) standard Hertz model and (b) the finite - thickness corrected Hertz model. (c) Young's modulus median value of single cell clusters ($E_{md}$, see Materials and Methods). (d-e) median Young's modulus ± $\sigma_E$; n = 3 clusters per condition. (f) mean Young's modulus ± standard error; n = 76-94 cells per condition.*

To verify that the results of the mechanical analysis are not affected by the change in the height of the cells inside the clusters (for example, the thickening of cells could determine a loss of sensitivity to the inner region of the cells, for the same maximum indentation, or to a redistribution of organelles in the cytoplasmic region), we plotted the median of Young's modulus from the inner cells body (nuclear region, deep indentation) of each cluster versus the mean cluster height (**Suppl. Fig. S4c**). There is no evidence of any correlation between the height of the cluster or the cell and Young's modulus, which led us to exclude that our analysis is significantly biased by the morphological changes of the cells. It is clear from the comparison of **Suppl. Fig. S4a and S4b** that the measured Young's modulus increases as the cell height decreases, leading to severe overestimation of the intrinsic (effective) value of a cell modulus. Noticeably, the substrate-induced also in the highest points. In our experimental conditions, only with cell heights of approximately 20-30 μm would the finite-thickness effect negligible.

The effect of the rigid substrate underneath the cells is to hide the fine mechanical modifications that take place inside the cells body, upon the action of the drug, resulting in a loss of sensitivity. Since we have shown that the morphological properties, and in particular the height of the cells, change upon the treatment with the drugs, the application of the finite-thickness correction is important to compare results obtained at different drug concentrations. This is shown in **Suppl. Fig. S4d-f**, where the results obtained with VFL with and without the finite-thickness correction are shown. Without the correction, apart from a rigid shift to higher Young's modulus, the differences between the three conditions are lost; with the application of the correction, instead, the differences are evident and the same trend is observed as with the pyramidal probes.

#### ***Supplementary Note 6: 3D organization of MTs visualized using confocal microscopy***

To visualize the 3D organization of MTs, movies were prepared by combining all recorded z-stacks for DU145 cells treated with low MTDs doses. List of movies:

- DU145_MTs_NTC.mp4
- DU145_MTs_VFL.mp4

- DU145_MTs_DTX.mp4
- DU145_MTs_COL.mp4

***Supplementary Note 7: Actin organization inside DU145 cells***

Maximum fluorescence intensity (maximum intensity projections of the Z-stacks of microscope images) obtained from phalloidin conjugated with Alexa Fluor 488 showing the organization of actin filaments in non-treated (NTC) and MTD–treated prostate DU145 cancer cells (**Suppl. Fig. S5)**.

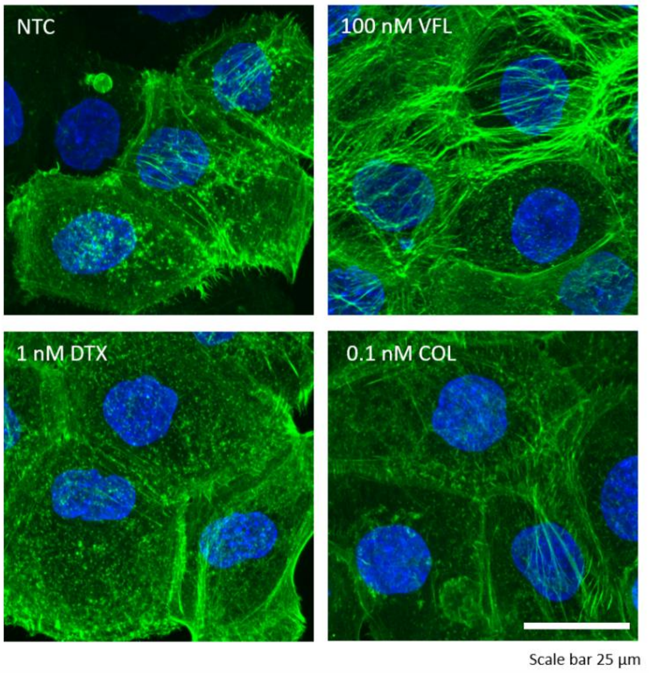


***Suppl. Figure S5****. Organization of actin filaments in non-treated (NTC) and MTDs-treated prostate DU145 cancer cells visualized using confocal microscopy.*

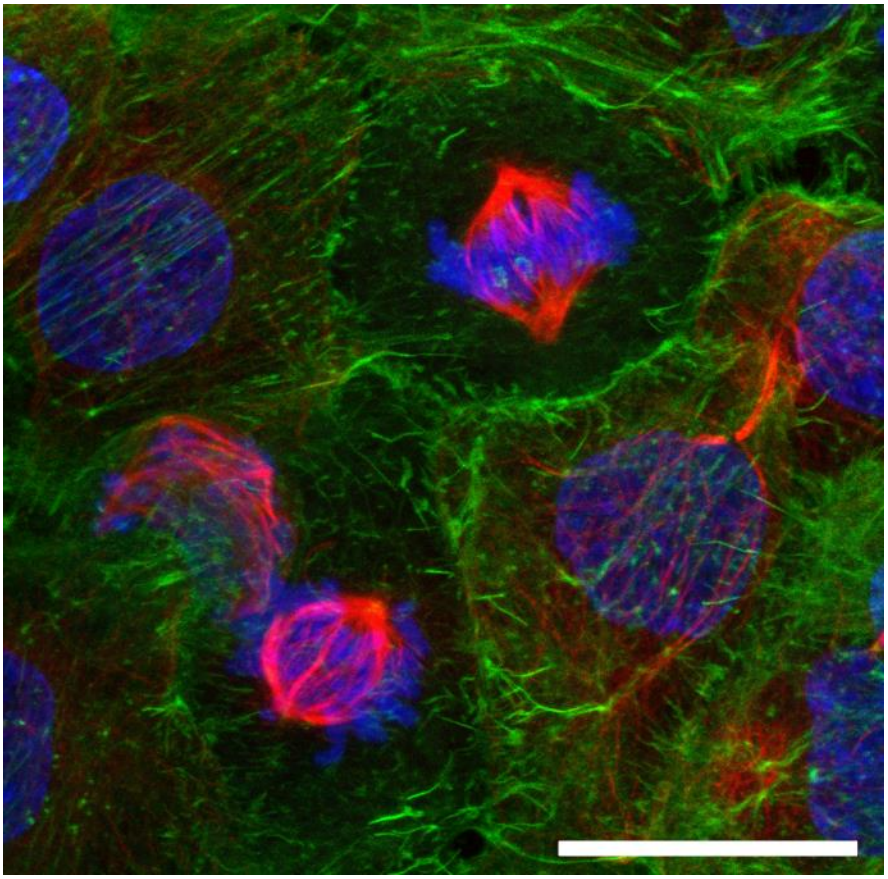

***Suppl. Figure S6****. Example of proliferating DU145 cells during the mitosis process after 72h of treatment with VFL. Maximal projection image from the fluorescent confocal microscope. Scale bar = 25 μm.*
